# Quenching of Chaos in externally driven metacommunities

**DOI:** 10.1101/2025.08.21.671425

**Authors:** Anusree Vinod Kunnath, Dweepabiswa Bagchi

## Abstract

Population dynamics of different species across food web models of diverse spatial scales have been extensively investigated over the past three decades. Chaotic fluctuations in species populations are generally associated with increased extinction risk due to frequent periods of low species density leading to cascading effects. The existence of a food web where the cohabiting species population oscillates chaotically has been largely attributed to chaos control or death of chaos, mediated either by internal mechanisms, such as the food web favoring parameter values that keep the species population out of the chaotic region, or external mechanisms like environmental forcing. Yet, this picture is not complete, as food webs do not exist in isolation, but are generally connected to each other through dispersal of species, forming a metacommunity. A metacommunity of food webs whose dynamics is that of chaotic attractors can only persist through control and particularly the the quenching of chaos. We claim that habitat heterogeneity in such a metacommunity fulfills this purpose. We address this question by analyzing the dynamics of a drive-response metacommunity composed of five chaotic food webs, each located in a distinct patch. Each patch contains an inherently chaotic tritrophic food web, with habitat heterogeneity present among patches. The first patch functions as the drive, exerting external influence on the dynamics of the remaining four response patches. Our results establish that in a drive-response metacommunity, strong influence from the drive quenches dynamical chaos in both drive and response patches, often leading to steady states. This phenomenon is observed in two distinct metacommunity network structures. The heterogeneity of the response systems (food web models) and the dissimilarity between drive and response systems are found to play significant roles in suppressing chaotic population dynamics. These findings strongly imply that the persistence of such inherently chaotic metacommunities may result from the quenching of chaos through the interplay of chaos, habitat heterogeneity, and to some degree network structure shaped by dispersal. Furthermore, we illustrate that these metacommunities are also susceptible to extinction due to dispersal-induced synchronization. Accordingly, this study also investigates dispersal-induced complete synchronization among the constituent patches.

## 1. Introduction

Chaotic population dynamics have been reported in several food web models describing the evolution of a plethora of natural ecosystems (Hastings and Powell (1991); Blasius et al. (1999); Upadhyay et al. (1998)). Most importantly, the extinction risk and the persistence of such inherently chaotic food webs have also been investigated in detail. Several studies have revealed that food web models in dynamically chaotic states have a high probability of suffering species extinction, either through cascading effects or a synchronized response to external environmental noise (Williams and Martineza (2004); Boccaletti et al. (2000)). In particular, Berryman and MillStein analyzed the phase space of several chaotic food web models and reported that low-density populations have an elevated chance of extinction in the chaotic domain of the phase space (Berryman and Millstein (1989)). This arises due to population extinctions under stochasticity (Thomas et al. (1980); Hastings (1982); Croteau (2010)). Additionally, chaotic dynamics might mask early warning signals and thereby indirectly increase extinction risk (Krkošek and Drake (2014)) of species. Further, Dietze (Dietze (2018)) reported that truly chaotic populations fluctuate wildly, resulting in frequent population crashes, genetic bottlenecks, and a high risk of stochastic extinction.

Speaking from a dynamical analysis perspective, several studies (Jørgensen and Jørgensen (2002); Berryman and Millstein (1989); Doebeli (1993)) have asserted that natural selection disfavors parameter regimes that would induce chaotic population dynamics of any species. Additionally, ecological mechanisms such as toxin release (Mandal et al. (2006)) and omnivory (McCann and Hastings (1997)) have been reported to regulate or suppress chaos in the population dynamics portrayed by several food web population models (Doebeli (1993); Gamarra et al. (2001)).

However, food webs rarely exist in isolation and are typically embedded within dispersally connected networks. Despite this, the suppression or quenching of chaos in such metacommunities has only been investigated to a limited extent. In case of such metacommunities, environmental factors like habitat heterogeneity (Krkošek and Drake (2014); Hilborn et al. (2003)) have been reported to supress chaotic behavior of population dynamics. In contrast, Allen et. al. (Allen et al. (1993)) found that chaos in a weakly coupled metapopulation can have a decorrelating effect, thereby enhancing persistence of the metacommunity. Speaking from a dynamical perspective, Solé and Gamarra (Solé and Gamarra (1998)) analyzed the phase space of a network of chaotic food web models and showed that it consists of an extensive chaotic regime. Within this domain, some dynamical regions lead to the metapopulation exhibiting chaotic population fluctuations that increase extinction risk, while other in other regimes desynchronization induced by chaos reduces the probability of global extinction. Most importantly, strong dispersal between the chaotic food webs can increase the extinction risk of the metapopulation by shifting their dynamics into the chaotic regime characterized by low fluctuations and high extinction risk (Solé and Gamarra (1998)). These findings strongly suggest that whether the existence of chaotic behavior in a metapopulation dynamics reduces their persistence or not depends on the strength of dispersal between the food webs. Thus, the influence of dispersal on the persistence of metacommunities with chaotic population dynamics warrants further investigation.

The structure of a metacommunity also plays a role in determining the persistence of the metapopulation. Although directional coupling among populations and food webs is well documented. (Leibold et al. (2004); Mouquet and Loreau (2003)), metacommunities explicitly structured as drive–response networks have received comparatively little attention. Indeed, many real metacommunities are structured by drive-response coupling, in the forms of asymmetric or hierarchical dispersal, as seen in riverine systems, marine currents, and source–sink landscapes (Amarasekare (2008); Holt (1997)).

From the perspective of dynamical analysis, drive–response networks reconcile the coexistence of stability and chaos, and their role in persistence of large-scale metacommunities (Pecora and Carroll (1990); Goldwyn and Hastings (2009)). Further, the importance of drive-response metacommunities in determining early signals of extinction risk is also paramount Bagchi et al. (2024).

Motivated by the above discussions, this study examines the quenching of chaos in a drive-response metacommunity. The metacommunity model consists of five patches, each characterized by inherently chaotic food webs that describe the species dynamics within each patch. The structure we consider for our metacommunity is that of a drive-response network, for e.g., a river-distributary ecosystem. In such a network, the river controls the water level, nutrient level, and soil level of the distributaries (Larsen et al. (2021)), thus effectively controlling their habitat quality (Sarremejane et al. (2021); J. D. Yeakel and deAguiar (2014)) and ‘driving’ their dynamics. To ensure that our model adheres to real metacommunities, we consider an appreciable degree of heterogeneity between drive and response patches. We demonstrate how habitat heterogeneity in such metacommunities quenches the chaos in each patch. Further, we establish that the complete synchronization between homogeneous drive and response systems is absolute, but decreases considerably with increasing dispersal in the case of heterogeneous drive and response. We also propose that our model could also be used as an approach to understand the effect of invasion in impeding chaos in the metacommunity.

## 2. Models

In our study, both the drive and response patches contain of a tri-trophic, inherently chaotic foodweb described by the Blasius model (Blasius et al. (1999)). The basic equations of the Blasius model is given as

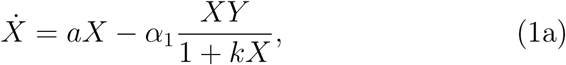

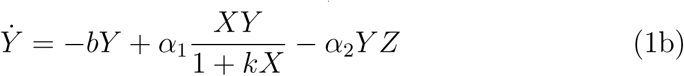

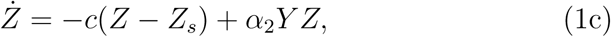

where the parameter *a* represents the growth rate of the resource in absence of inter-species interactions, while the parameters *b* and *c* represnts the mortality rate of the predator and consumer. The parameter *α*_1_ and *α*_2_ represents the predation rate of the predator and that of the consumer respectively, while the predator mortality rate is given by *b*.

The food web of the drive patch is described by the Blasius model (Appendix A). The predator mortality rate of the drive patch is given by *b_d_*. The amplitude of drive dispersal is given by *e*. The predator mortality rate of the drive patch is given by *b_d_*. The amplitude of drive dispersal is given by *e*. Equivalently, the food web of the response patch is governed by the same Blasius Model, with a different dispersal term as explained in Appendix B. For the *i^th^* response patch, the parameter *b_i_* describes the mortality of the predator of the *i^th^* patch. The other parameters are same as above.

## 3. Results

Figure 1 shows the bifurcation diagram of the Blasius model food web with respect to the mortality of predator *b*. We have chosen both *b_d_* and all *b_r_* values such that our investigated food webs all exhibit chaotic dynamics, when unperturbed (*b ɛ* (0.8 − 1)). The structure of our metacommunity is represented in 1(b), where the circle filled with red represents the drive patch, the arrows from it represent the drive-response diseprsal. The circles filled with green represent the response patches. The dotted lines represent interresponse dispersals, whose magnitude is 0 when we consider unconnected responses. As stated above, the habitat heterogeneity across various patches in the metacommnunity comes from the variation in predator mortality rate *b* from one patch to another.

**Figure 1:**
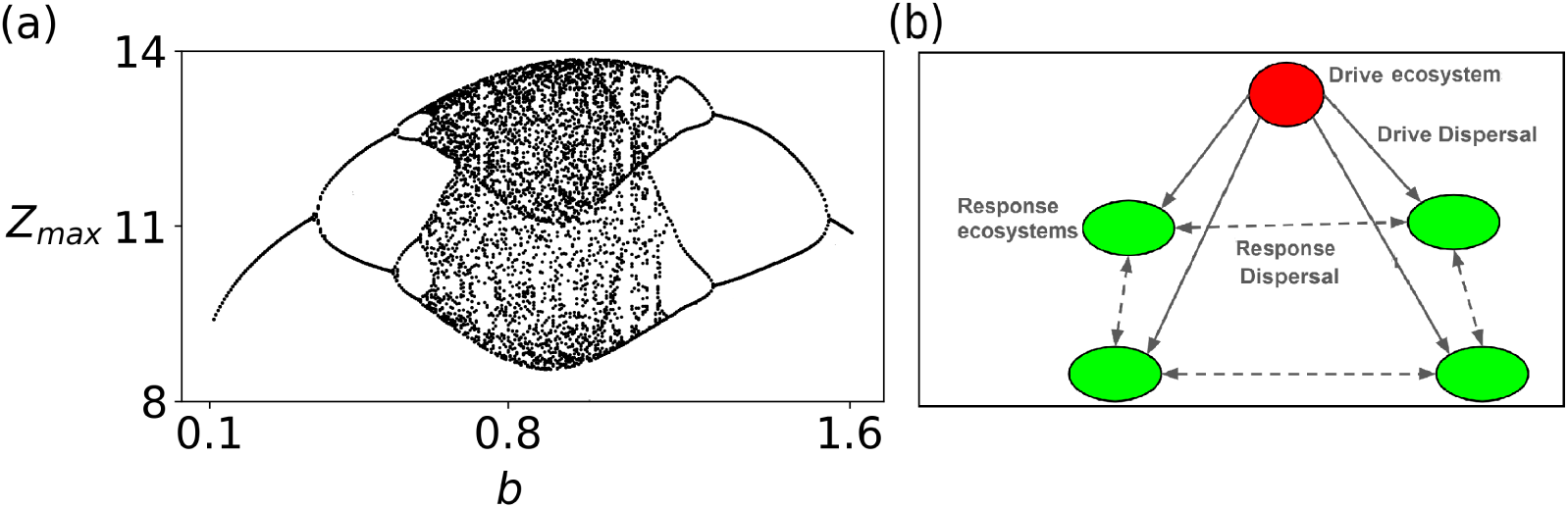
(a) Bifurcation diagram of one single food web model described by the Blasius model, plotted as a function of predator mortality rate *b*.

### 3.1. Locally connected response patches

#### 3.1.1. Homogeneous drive and response patches

First, we consider a group of locally connected, completely identical responses that are linked to an identical drive through dispersal, where we have fixed the predator mortality rate for both drive and response patches as *b_drive_* = *b_responses_* = 0.9. The temporal dynamics of the consumers from the drive patch *Z_d_* and a representative response patch *Z_r_* are depicted in Fig. 2(a) and 2(b) for the dispersal strengths *e* = 0.2 and 3.5, respectively. The chaotic attractor of the both drive patch (red line) and a representative response patch (green line) are also shown in the inset of both Figs. 2(a) and 2(b). It is evident from these figures that the time series of *Z_d_* and *Z_r_* evolve identically corroborating the complete synchronization among the drive and uncoupled response systems for both low and high values of the dispersal strength. In order to elucidate the nature of the dynamics exhibited by both drive and response systems, we have plotted their bifurcation diagrams, obtained by collecting the maxima *Z_dmax_* and *Z_rmax_* of the temporal dynamics of the drive and response patches, in Figs. 2(c) and 2(d), respectively, in the range of the dispersal strength *e* ∈ (0.0, 3.5). It is evident from these bifurcation diagrams that the evolution of the consumer, and consequently resource *X* and predator *Y*, of both drive and response patches exhibit chaotic oscillation for their population density in the entire explored range of the dispersal strength. Thus, the homogeneous drive is not capable of quenching the chaotic evolution of the population density in response patches.

**Figure 2:**
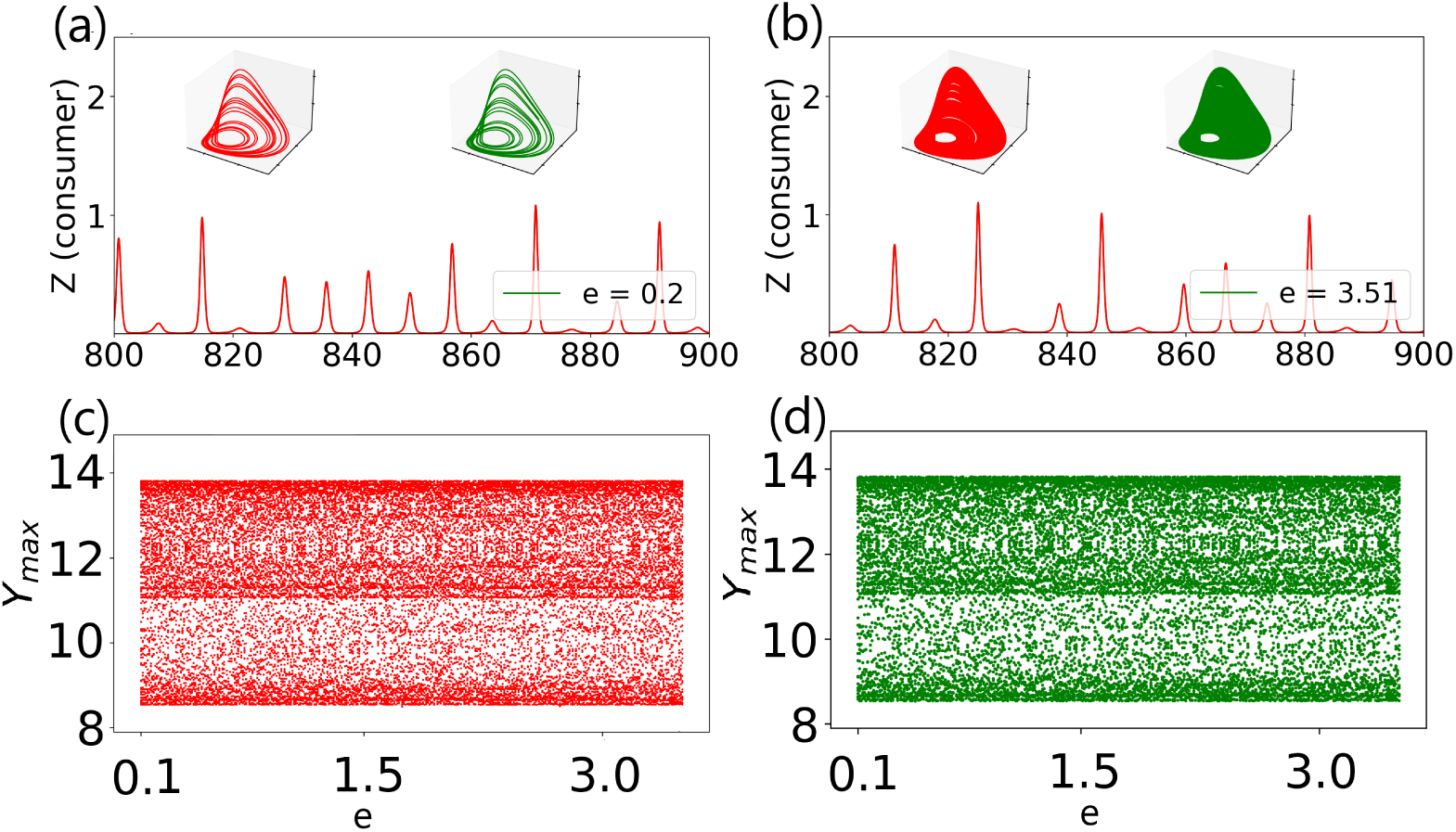
Locally connected response patches: Completely identical drive and homogeneous response patches. (a) Time series of consumer dynamics of both drive patch and a response patch for the low dispersal strength *e* = 0.2, (b) same as (a) for the high dispersal strength *e* = 3.5, (c) bifurcation diagram of the drive patch as a function of *e*, and (d) bifurcation diagram of a response patch as a function of *e*. Attractors of the drive (red line) and a response (green line) patches are depicted in the inset of (a) and (b). Predator mortality rate for drive *b_d_*= 0.9, and for responses *b_r_* = 0.8.

#### 3.1.2. Heterogeneous drive and homogeneous response patches

Next, we consider a group of locally connected, completely identical responses that are linked to a non-identical drive through dispersal, where we have fixed the predator mortality rate for both drive and response patches as *b_drive_* = 0.9 and *b_responses_*= 0.8. The temporal dynamics of the consumers from the drive patch *Z_d_* and a representative response patch *Z_r_* are depicted in Fig. 3(a) and 3(b) for the dispersal strengths *e* = 0.2 and 3.5, respectively. The chaotic attractor of the both drive patch (red line) and a representative response patch (green line) are also shown in the inset of both Figs. 3(a) and 3(b). In order to elucidate the nature of the dynamics exhibited by both drive and response systems, we have plotted their bifurcation diagrams, obtained by collecting the maxima *Z_dmax_* and *Z_rmax_* of the temporal dynamics of the drive and response patches, in Figs. 3(c) and 3(d), respectively, in the range of the dispersal strength *e* ∈ (0.0, 3.5). It is evident from these bifurcation diagrams that the evolution of the consumer, and consequently resource *X* and predator *Y*, of both drive and response patches exhibit chaotic oscillation for their population density in the range of low dispersal strength. However, with increasing dispersal (*eɛ*(0.8, 1.0)), the dynamics of the drive patche undergo Hopf bifurcation from chaotic to a bistable state, consisting of 2 limit cycles. Upon further increase of dispersal (*eɛ*(1.25, 3.5)) the dynamics of the drive patch further undergoes reverse bifurcation to obtain a bistable steady state. However, in case of the response patch, with increasing dispersal (*eɛ*(0.8, 1.0)) the dynamics undergoes Hopf Bifurcation from chaotic go mutli-stable limit cycles. Upon further increase of dispersal (*eɛ*(1.25, 3.5)) the dynamics of the drive patch further undergoes reverse bifurcation to obtain a multi-stable steady state. Thus, the on-identical but similar drive quenches the chaotic evolution of the population density in completely identical locally connected response patches.

**Figure 3:**
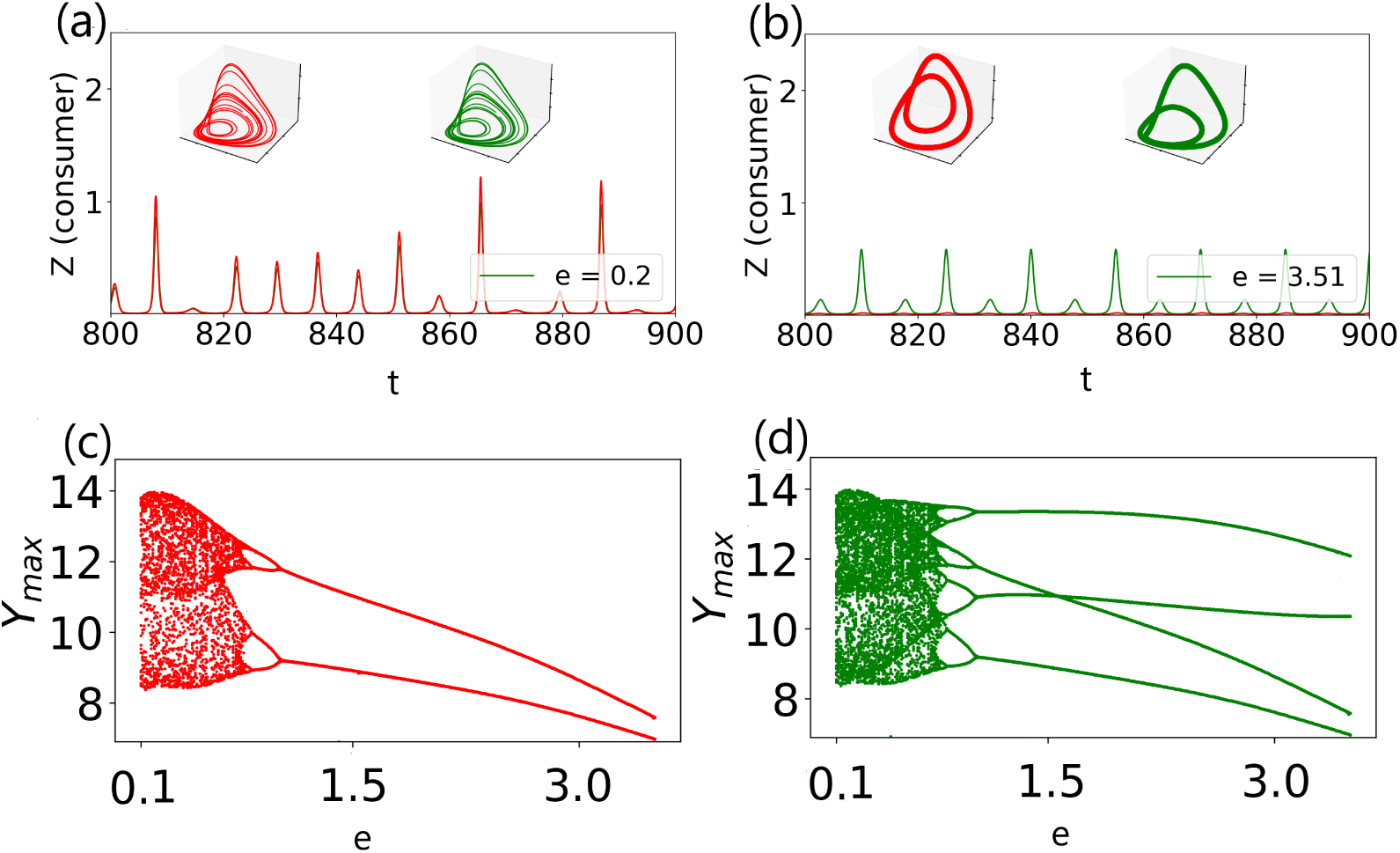
Locally connected response patches: Nonidentical drive and homogeneous response patches. (a) Time series of consumer dynamics of both drive patch and a response patch for the low dispersal strength *e* = 0.2, (b) same as (a) for the high dispersal strength *ε_d_* = 3.5, (c) bifurcation diagram of the drive patch as a function of *e*, and (d) bifurcation diagram of a response patch as a function of *e*. Attractors of the drive (red line) and a response (green line) patches are depicted in the inset of (a) and (b). Predator mortality rate for drive *b_d_* = 0.9, and for responses *b_r_* = 0.8.

#### 3.1.3. Heterogeneous drive and response patches

Next, we consider a group of locally connected, heterogeneous responses that are linked to a non-identical drive through dispersal, where we have fixed the predator mortality rate for both drive and response patches as *b_drive_* = 0.9 and *b_responses_*= 0.77, 0.76,0.82 and 0.85. The temporal dynamics of the consumers from the drive patch *Z_d_* and a representative response patch *Z_r_* are depicted in Fig. 4(a) and 4(b) for the dispersal strengths *e* = 0.2 and 3.5, respectively. The dynamics evidently changes from chaotic to steady state on increasing drive-response dispersal. The chaotic attractor of the both drive patch (red line) and a representative response patch (green line) are also shown in the inset of both Figs. 4(a) and 4(b). In order to elucidate the nature of the dynamics exhibited by both drive and response systems, we have plotted their bifurcation diagrams, obtained by collecting the maxima *Z_dmax_* and *Z_rmax_* of the temporal dynamics of the drive and response patches, in Figs. 4(c) and 4(d), respectively, in the range of the dispersal strength *ε_d_* ∈ (0.0, 3.0). It is evident from these bifurcation diagrams that the evolution of the consumer, and consequently resource *X* and predator *Y*, of both drive and response patches exhibit chaotic oscillation for their population density in the range of low dispersal strength (*eϵ*(0.1, 0.25)). However, with increasing dispersal (*eϵ*(0.3, 0.8)), the dynamics of the drive patch undergo Hopf bifurcation from chaotic to a bistable state, consisting of 2 limit cycles. Upon further increase of dispersal (*eϵ*(0.8, 2.5)) the dynamics of the drive patch further undergoes reverse bifurcation to obtain a monostable steady state, with a decreasing trend in the average population density. On further increase in dispersal, (*e >* 2.5), the mono stable state is maintained but the average population density saturates. In case of the response patch, with increasing dispersal (*eϵ*(0.1, 0.25)) the dynamics stays chaotic. Upon further increase of dispersal (*eϵ*(0.25, 0.8)) the dynamics of the drive patch further undergoes reverse bifurcation to obtain a multi-stable limit cycle. Upon further increase of dispersal (*eϵ*(0.8, 2.5)) the dynamics of the drive patch further undergoes reverse bifurcation to obtain a bistable steady state, with a decreasing trend in the average population density. On further increase in dispersal, (*e >* 2.5), the bistable state is maintained but the average population density saturates. Thus, the non-identical but similar drive quenches the chaotic evolution of the population density in non-identical locally connected response patches.

**Figure 4:**
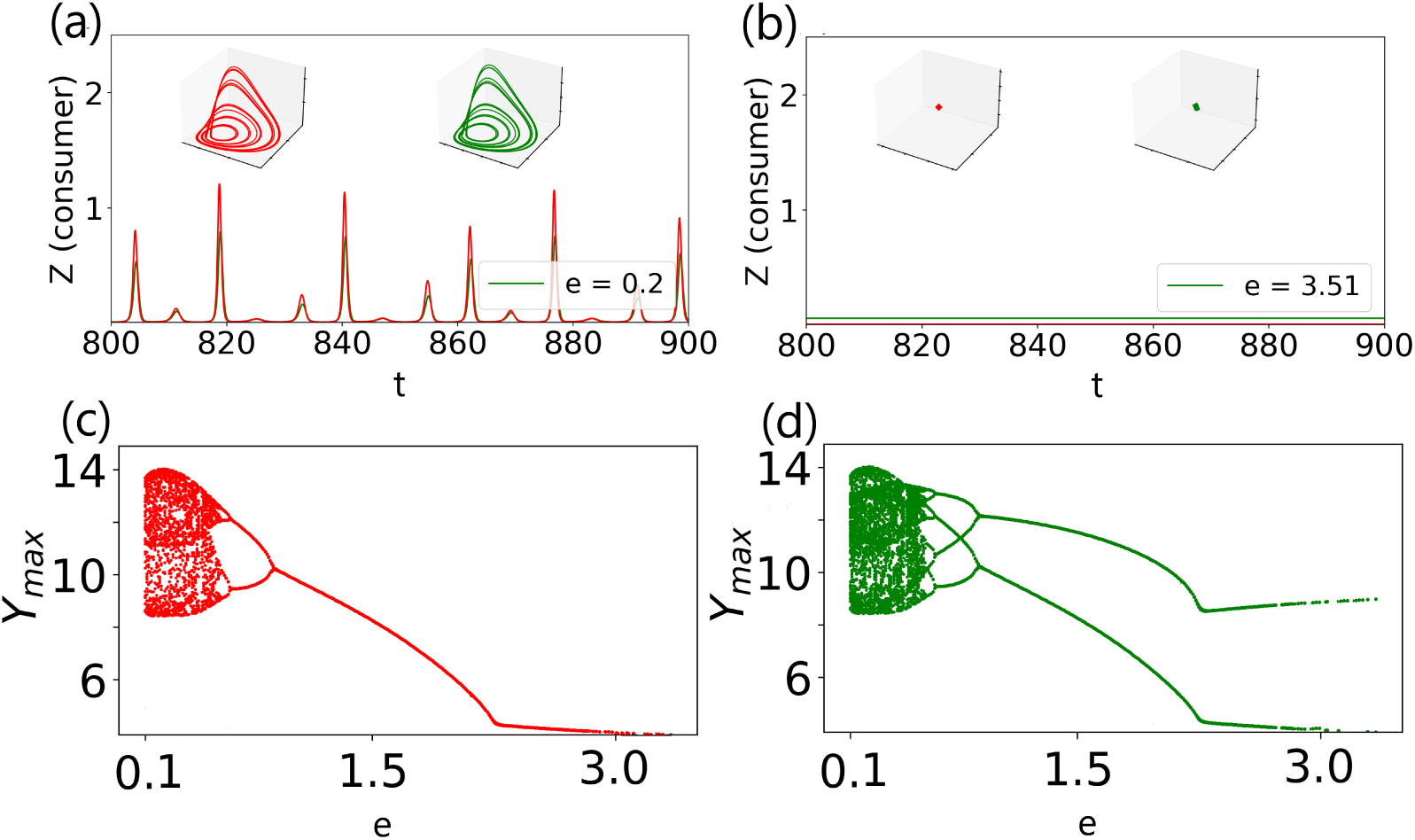
Same as Fig. 3 but predator mortality rate for drive and response patches are *b_d_* = 0.9 and *b_r_* = 0.77, 0.76,0.82 and 0.85.

**Figure 5:**
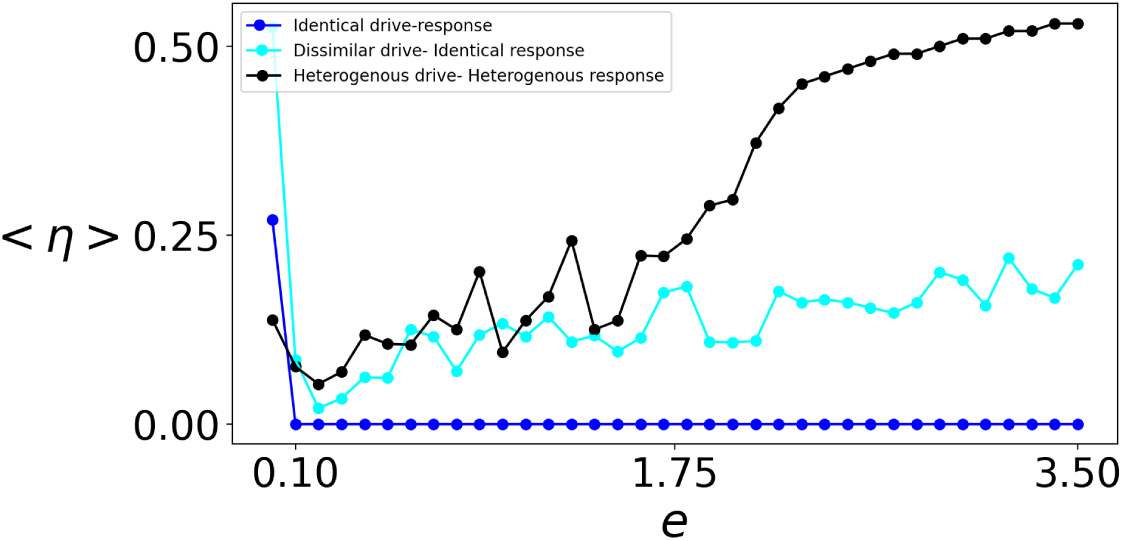
N

#### 3.1.4. Synchronization of the metacommunity

We investigate the emergence of synchronization in a group of oscillators as a function of coupling strength, considering identical, dissimilar and heterogenous systems. Our goal is to determine what degree of heterogeneity promotoes synchrony and what level of heterogeneity is required in the network to induce asynchronous steady states. We define synchronization error for the group of oscillators. averaged over time as,

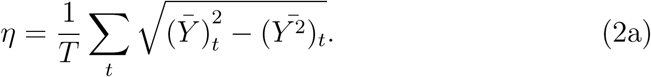

Here, 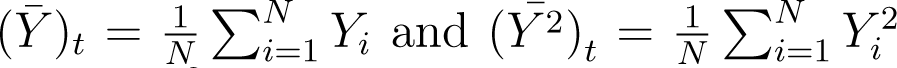 average values of the variables *Y* and *Y*^2^ of oscillators *i* = 1, *…N*, at an instant of time *t*. Further, we average *η* over different initial states to obtain an ensemble averaged synchronization error *η*.

Figure 5 demonstrates the synchronization between all the patches in the metacommunity, when the responses are nearest neighbor coupled. When the drive and responses are completely identical, all patches are synchronized. However, with slight habitat heterogeneity in the system, the synchronization error has a low but non-zero magnitude. However, with increasing invasionary dispersal *e*, the synchronization between the patches decrease. Synchronization is detrimental to the persistence of the metacommunity. Therefore, it is clearly demonstrated that in this case, invasion increases the persistence of the metacommunity as a whole. With further increase in the heterogeneity in the network, for the range of dispersal *eɛ*(0.1 − 1.5, there is an increasing but fluctuating trend in the synchronization error *< η >*. However, with further increase in dispersal *e*, the synchronization error keeps steadily increasing without fluctuation.

### 3.2. Globally connected response patches

#### 3.2.1. Homogeneous drive and response patches

First, we consider a group of globally connected, completely identical responses that are linked to an identical drive through dispersal, where we have fixed the predator mortality rate for both drive and response patches as *b_drive_* = *b_responses_* = 0.9. The temporal dynamics of the consumers from the drive patch *Z_d_* and a representative response patch *Z_r_* are depicted in Fig. 6(a) and 6(b) for the dispersal strengths *e* = 0.2 and 3.5, respectively. The chaotic attractor of the both drive patch (red line) and a representative response patch (green line) are also shown in the inset of both Figs. 6(a) and 6(b). It is evident from these figures that the time series of *Z_d_* and *Z_r_* evolve identically corroborating the complete synchronization among the drive and uncoupled response systems for both low and high values of the dispersal strength. In order to elucidate the nature of the dynamics exhibited by both drive and response systems, we have plotted their bifurcation diagrams, obtained by collecting the maxima *Z_dmax_* and *Z_rmax_* of the temporal dynamics of the drive and response patches, in Figs. 6(c) and 6(d), respectively, in the range of the dispersal strength *e* ∈ (0.0, 3.5). It is evident from these bifurcation diagrams that the evolution of the consumer, and consequently resource *X* and predator *Y*, of both drive and response patches exhibit chaotic oscillation for their population density in the entire explored range of the dispersal strength. Thus, the homogeneous drive is not capable of quenching the chaotic evolution of the population density in response patches.

**Figure 6:**
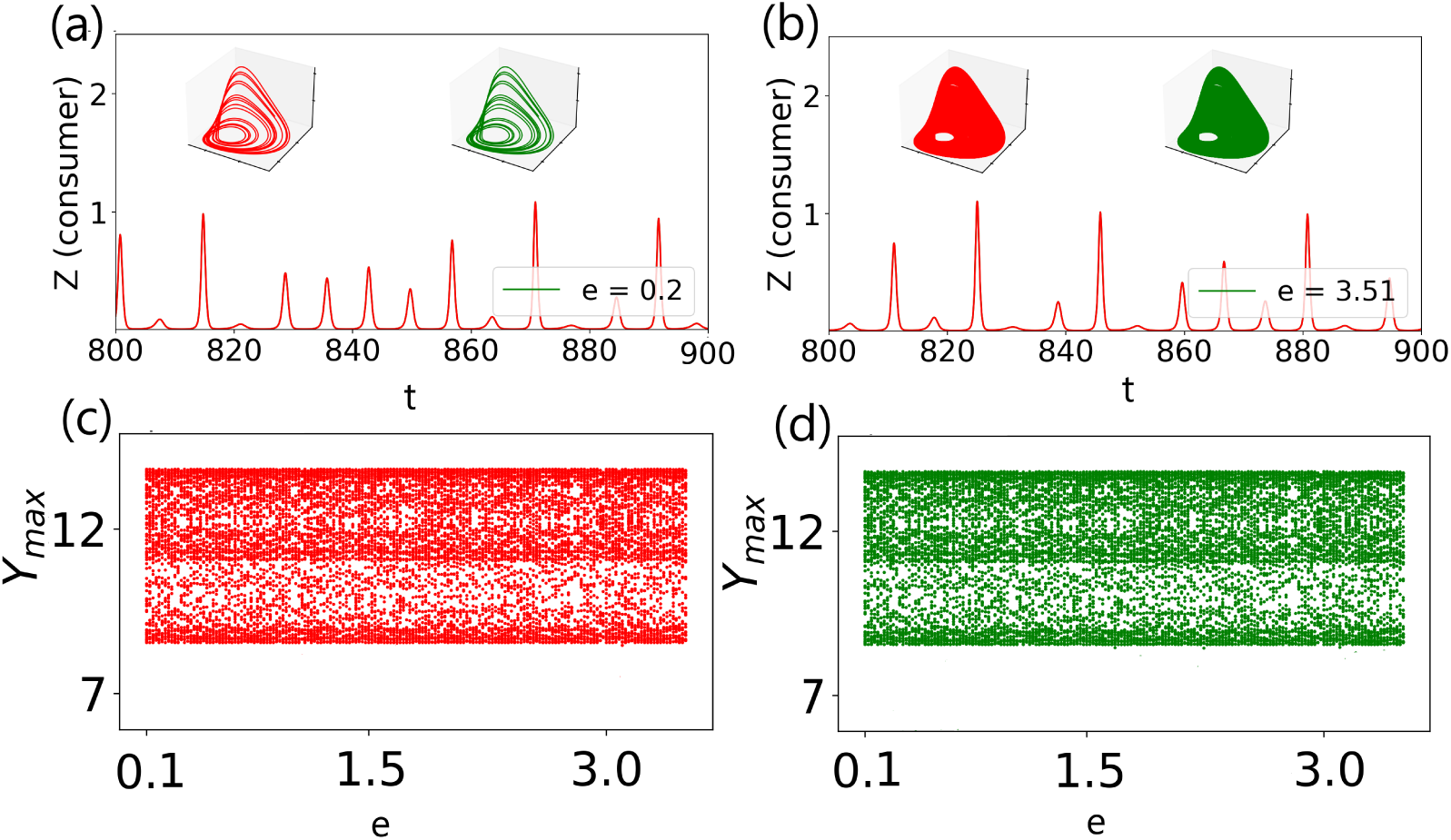
Globally connected response patches: Completely identical drive and homogeneous response patches. (a) Time series of consumer dynamics of both drive patch and a response patch for the low dispersal strength *e* = 0.2, (b) same as (a) for the high dispersal strength *ε_d_* = 3.5, (c) bifurcation diagram of the drive patch as a function of *e*, and (d) bifurcation diagram of a response patch as a function of *e*. Attractors of the drive (red line) and a response (green line) patches are depicted in the inset of (a) and (b). Predator mortality rate for drive *b_d_*= 0.9, and for responses *b_r_* = 0.8.

#### 3.2.2. Heterogeneous drive and homogeneous response patches

Next, we consider a group of globally connected, completely identical responses that are linked to a non-identical drive through dispersal, where we have fixed the predator mortality rate for both drive and response patches as *b_drive_*= 0.9 and *b_responses_* = 0.8. The temporal dynamics of the consumers from the drive patch *Z_d_* and a representative response patch *Z_r_* are depicted in Fig. 7(a) and 7(b) for the dispersal strengths *e* = 0.2 and 3.5, respectively. The chaotic attractor of the both drive patch (red line) and a representative response patch (green line) are also shown in the inset of both Figs. 7(a) and 7(b). In order to elucidate the nature of the dynamics exhibited by both drive and response systems, we have plotted their bifurcation diagrams, obtained by collecting the maxima *Z_dmax_* and *Z_rmax_* of the temporal dynamics of the drive and response patches, in Figs. 7(c) and 7(d), respectively, in the range of the dispersal strength *ε_d_*∈ (0.0, 3.5). It is evident from these bifurcation diagrams that the evolution of the consumer, and consequently resource *X* and predator *Y*, of both drive and response patches exhibit chaotic oscillation for their population density in the range of low dispersal strength. However, with increasing dispersal (*eɛ*(0.8, 1.0)), the dynamics of the drive patche undergo Hopf bifurcation from chaotic to a bistable state, consisting of 2 limit cycles. Upon further increase of dispersal (*eɛ*(1.25, 3.5)) the dynamics of the drive patch further undergoes reverse bifurcation to obtain a bistable steady state. However, in case of the response patch, with increasing dispersal (*eɛ*(0.8, 1.0)) the dynamics undergoes Hopf Bifurcation from chaotic go mutli-stable limit cycles. Upon further increase of dispersal (*eɛ*(1.25, 3.5)) the dynamics of the drive patch further undergoes reverse bifurcation to obtain a multi-stable steady state. Thus, the on-identical but similar drive quenches the chaotic evolution of the population density in completely identical locally connected response patches.

**Figure 7:**
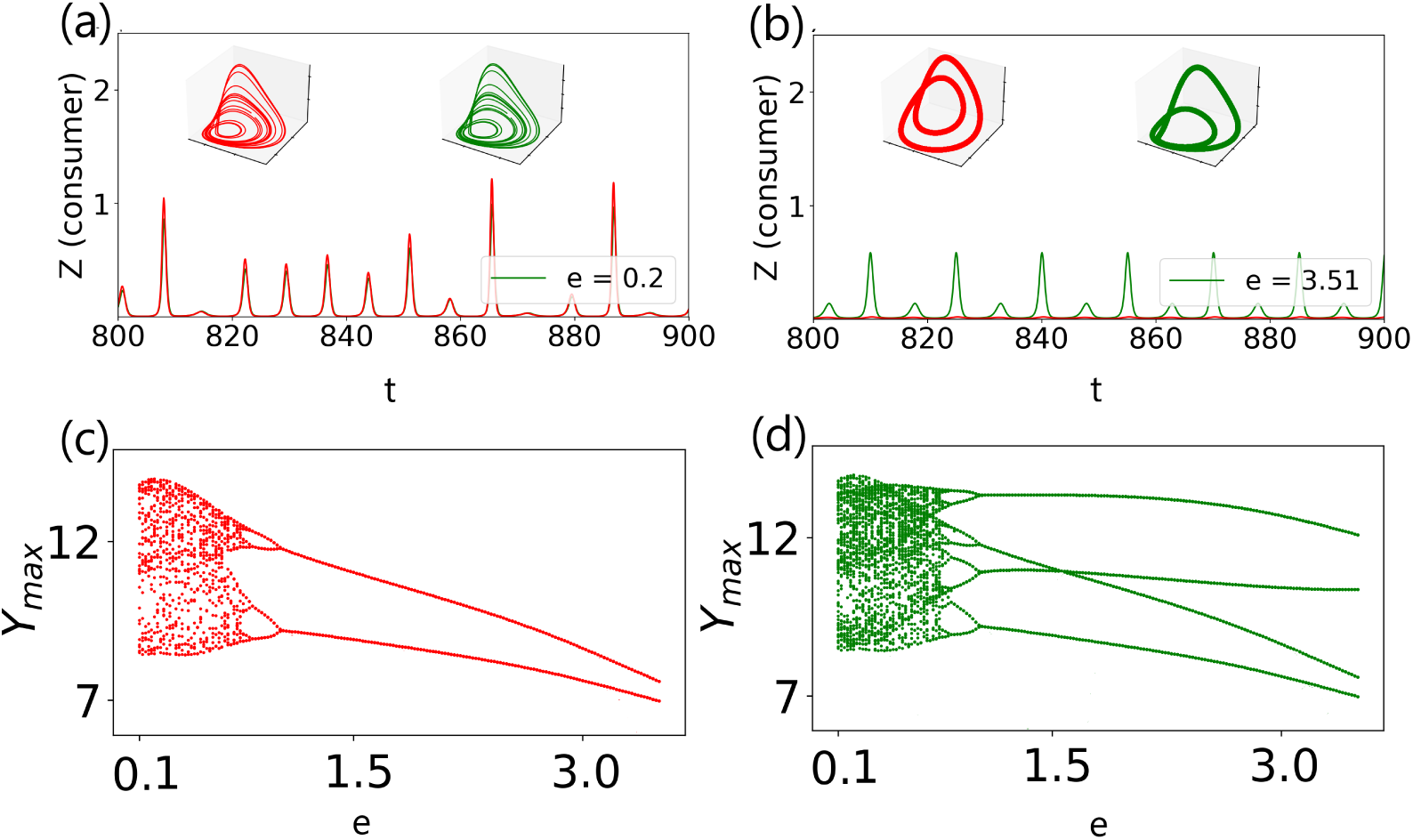
Same as Fig. 6 but predator mortality rate for drive and response patches are *b_d_* = 0.9 and *b_r_* = 0.8.

#### 3.2.3. Heterogeneous drive and homogeneous response patches

Next, we consider a group of locally connected, heterogeneous responses that are linked to a non-identical drive through dispersal, where we have fixed the predator mortality rate for both drive and response patches as *b_drive_* = 0.9 and *b_responses_*= 0.77, 0.76,0.82 and 0.85. The temporal dynamics of the consumers from the drive patch *Z_d_* and a representative response patch *Z_r_* are depicted in Fig. 8(a) and 8(b) for the dispersal strengths *e* = 0.2 and 3.5, respectively. The dynamics evidently changes from chaotic to steady state on increasing drive-response dispersal. The chaotic attractor of the both drive patch (red line) and a representative response patch (green line) are also shown in the inset of both Figs. 8(a) and 8(b). In order to elucidate the nature of the dynamics exhibited by both drive and response systems, we have plotted their bifurcation diagrams, obtained by collecting the maxima *Z_dmax_* and *Z_rmax_* of the temporal dynamics of the drive and response patches, in Figs. 8(c) and 8(d), respectively, in the range of the dispersal strength *ε_d_*∈ (0.0, 3.0). It is evident from these bifurcation diagrams that the evolution of the consumer, and consequently resource *X* and predator *Y*, of both drive and response patches exhibit chaotic oscillation for their population density in the range of low dispersal strength (*eϵ*(0.1, 0.25)). However, with increasing dispersal (*eϵ*(0.4, 0.9)), the dynamics of the drive patch undergo Hopf bifurcation from chaotic to a multistable state, consisting of 4 limit cycles to ultimately bistable states consisting of 2 limit cycles. Upon further increase of dispersal (*eϵ*(0.8, 2.5)) the dynamics of the drive patch further undergoes reverse bifurcation to obtain a monostable steady state, with a decreasing trend in the average population density. On further increase in dispersal, (*e >* 2.5), the mono stable state is maintained but the average population density saturates. In case of the response patch, with increasing dispersal (*eϵ*(0.1, 0.25)) the dynamics stays chaotic. Upon further increase of dispersal (*eϵ*(0.25, 0.8)) the dynamics of the drive patch further undergoes reverse bifurcation to obtain a multi-stable limit cycle. Upon further increase of dispersal (*eϵ*(0.8, 2.5)) the dynamics of the drive patch further undergoes reverse bifurcation to obtain a bistable steady state, with a decreasing trend in the average population density. On further increase in dispersal, (*e >* 2.5), the bistable state is maintained but the average population density saturates. Thus, the non-identical but similar drive quenches the chaotic evolution of the population density in non-identical locally connected response patches.

**Figure 8:**
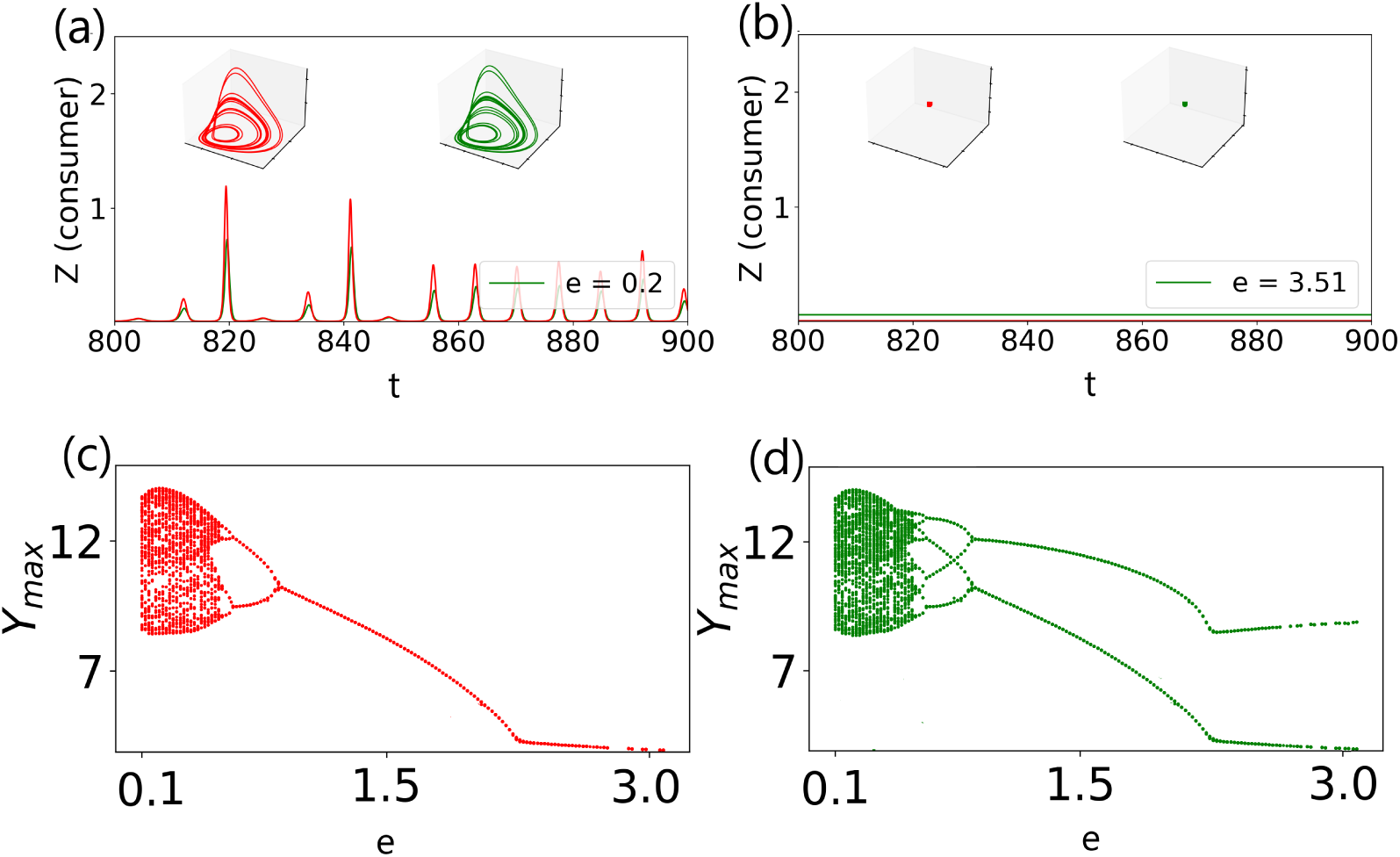
Same as Fig. 7 but predator mortality rate for drive and response patches are *b_d_* = 0.9 and *b_r_* = 0.77, 0.76,0.82 and 0.85.

**Figure 9:**
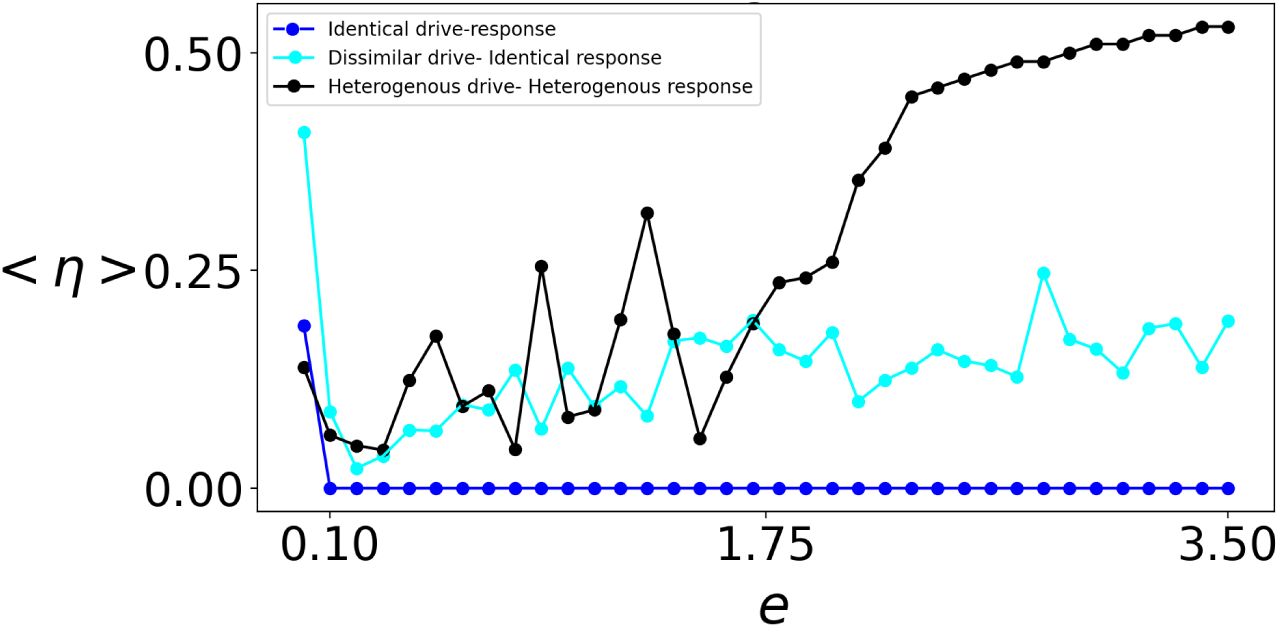
N

#### 3.2.4. Synchronization of the network

Figure 9 demonstrates the synchronization between the different patches of the metacommunity. The synchronizations follow the same nature as that of Fig. 5.

### 4. Discussion and Conclusion

Chaotic population dynamics in ecosystems, or specifically food web models have been studied in some detail. Population dynamics exhibiting chaotic behavior undergo rapid, large fluctuations. Such fluctuations subject populations to noise, genetic bottlenecks and extinctions induced by stochasticity. As a result, chaotic population dynamics have generally been associated with low persistence of food webs. However, ecosystems with inherently chaotic food webs continue to exist and thrive. Several studies suggest that this due to the control or quenching of chaos engineered by inherent ecological mechanisms or external forcings from the environment.

Even though there have been studies regarding the chaotic population dynamics of an inherently chaotic food web, dispersally connected networks of such food webs remain underexplored. Furthermore, existing studies show decidedly mixed outcomes. While some studies have reported that habitat heterogeneity can suppress chaotic population dynamics, others have established that weak dispersal between chaotic foodwebs can decorrelate populations and enhance metacommunity persistence. Phase space analyses of a network of chaotic food webs have been reported to consist of different chaotic regimes, which have opposing effects. While some chaotic regimes amplify fluctuations and extinction risk, others promote desynchronization that reduces global extinction. It has also been reported that in a metacommunity of chaotic foodwebs, strong dispersal can push the system into regimes of high extinction risk. These results indicate that the impact of chaos on meta-community persistence critically depends on dispersal strength, warranting further study.

Motivated by the above, we consider an ecological network of five patches, with each patch containing a tri-trophic, chaotic foodweb. We looked at how quenching of chaos occurs in both a locally connected and a globally connected network of ecosystems when driven by invasionary dispersal from an external ecosystem. For both types of network structures, we first considered the case of completely identical drive and responses. We demonstrated that for completely identical drives and responses, chaotic fluctuations are too strong to quench, irrespective of drive-response dispersal amplitude. Next, we introduced heterogeneity in predator mortality of the drive patch and response patches. This led to quenching of chaos and emergence of multistable steady states in the population dynamics of both drives and responses for high degrees of drive-response dispersal. Under a high degree of dispersal, the dynamics of the drive was that of a bistable state, and that of response was of a multi-stable steady state. Next, we increased the degree of heterogeneity even further and included heterogeneity in predator mortality across the response patches as well. As a result, the bistable steady state of the drive changed to a monostable steady state, and that of the responses changed to a bistable steady state. Our results strongly signify that heterogeneous meta-communities attain increased robustness and persistence through quenching of chaos.

Furthermore, for both kinds of network structure, we investigated various kinds of synchronizations induced by dispersal between the drive and response patches. Even with quenching of chaos, generalized synchronization is established in some, but not all, of the cases. The complete synchronization (quantified by synchronization error) and phase synchronization are nonexistent in most cases. These two types of synchronization are determined by network structure. However, the same investigation into synchronization between two response patches shows that in most cases, both phase and generalized synchronization exist. However, complete synchronization exists only when two responses are completely identical.

From the above results, we conclude that, barring a few idealized cases of completely identical drive and responses, dispersal primarily quenches chaos in real metacommunities consisting of inherently chaotic ecosystems and heterogeneous patches. Indeed, this study highlights the relevance of habitat heterogeneity in increasing persistence (or decreasing extinction risk) of metacommunities through control and quenching of chaos. This study also comments on the relevance of dispersal as one of the most fundamental mechanisms stabilizing the dynamics of metacommunities. Further, our work quantifies the level of drive dispersal required to quench chaos and whether such dispersal stabilizes the metacommunity or pushes it to extinction. Moreover, our investigation proves that whether the post-quenching steady-state dynamics has high species-density or low species-density depends on the degree of dispersal, habitat heterogeneity, and also on the network structure of the metacommunity.

## Appendix A. Appendix: Blasius Model (Drive System)

The food web of the drive patch is described by the Blasius model Blasius et al. (1999) and governed by the equations

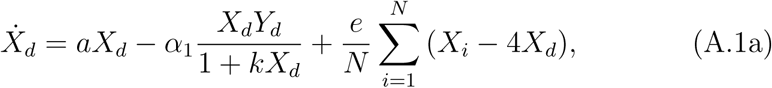

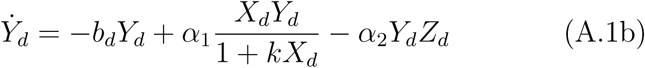

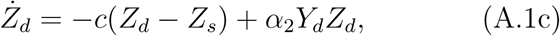

The parameter *a* represents the growth rate of the resource in absence of inter-species interactions, while the parameters *b* and *c* represnts the mortality rate of the predator and consumer. The parameter *α*_1_ represents the predation rate of the predator and *α*_2_ represents the predation rate of the consumer. The predator mortality rate of the drive patch is given by *b_d_*. The amplitude of drive dispersal is given by *e*. The parameter *N* represents the number of response patches. For this entire study, *N* = 4.

## Appendix B. Appendix: Blasius Model (Response network)

The food web of the response patches are governed by the following equations (Blasius Model):

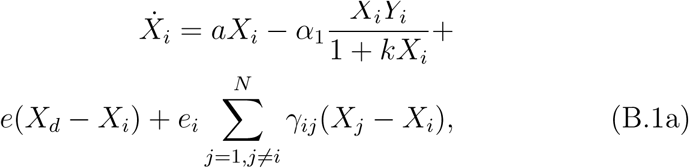

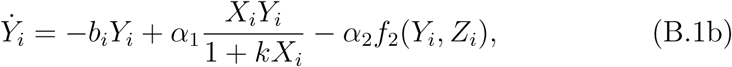

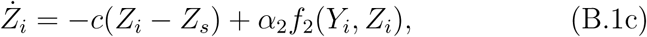

Γ*_ij_* is the adjacency matrix detailing the network structure created through dispersal between patches. When the response patches are unconnected, all elements of the matrix *γ_ij_ ɛ* Γ*_ij_* = 0. For nearest neighbor coupling between the responses, elements of the matrix *γ_ij_ ɛ* Γ*_ij_* ≠ 0, only if *i* = *j*. When the response patches are randomly coupled, elements of the matrix *γ_ij_ ɛ* Γ*_ij_* ≠ 0 randomly. For the response patch *i*, the parameter *b_i_* describes the mortality of the predator of the drive patch. The other parameters are given by *α*_1_ = 0.2, *α*_2_ = 1, *k* = 0.05, *Z_s_* = 0.006, *c* = 10.0, *a* = 1 and *f* = 0.25. The magnitude of simple diffusive dispersal between the *i^th^* resource patch and its nearest neighbors is given by the parameter *e_i_*. Note that when we consider the scenario for **unconnected** response patches *ε* = 0. For the *i^th^* response patch, the parameter *b_i_* describes the mortality of the predator of the *i^th^* patch. The other parameters are same as above.

## Acknowledgements

The work of D.B. is supported by IISER Thiruvananthapuram.

